# Development and pre-clinical characterization of two therapeutic equine formulations towards SARS-CoV-2 proteins for the potential treatment of COVID-19

**DOI:** 10.1101/2020.10.17.343863

**Authors:** Guillermo León, María Herrera, Mariángela Vargas, Mauricio Arguedas, Andrés Sánchez, Álvaro Segura, Aarón Gómez, Gabriela Solano, Eugenia Corrales-Aguilar, Kenneth Risner, Aarthi Narayanan, Charles Bailey, Mauren Villalta, Andrés Hernández, Adriana Sánchez, Daniel Cordero, Daniela Solano, Gina Durán, Eduardo Segura, Maykel Cerdas, Deibid Umaña, Edwin Moscoso, Ricardo Estrada, Jairo Gutiérrez, Marcos Méndez, Ana Cecilia Castillo, Laura Sánchez, José María Gutiérrez, Cecilia Díaz, Alberto Alape

## Abstract

In the current global emergency due to SARS-CoV-2 outbreak, passive immunotherapy emerges as a promising treatment for COVID-19. Among animal-derived products, equine formulations are still the cornerstone therapy for treating envenomations due to animal bites and stings. Therefore, drawing upon decades of experience in manufacturing snake antivenom, we developed and preclinically evaluated two anti-SARS-CoV-2 polyclonal equine formulations as potential alternative therapy for COVID-19. We immunized two groups of horses with either S1 (anti-S1) or a mixture of S1, N, and SEM mosaic (anti-Mix) viral recombinant proteins. Horses reached a maximum anti-viral antibody level at 7 weeks following priming, and showed no major adverse acute or chronic clinical alterations. Two whole-IgG formulations were prepared via hyperimmune plasma precipitation with caprylic acid and then formulated for parenteral use. Both preparations had similar physicochemical and microbiological quality and showed ELISA immunoreactivity towards S1 protein and the receptor binding domain (RBD). The anti-Mix formulation also presented immunoreactivity against N protein. Due to high anti-S1 and anti-RBD antibody content, final products exhibited high *in vitro* neutralizing capacity of SARS-CoV-2 infection, 80 times higher than a pool of human convalescent plasma. Pre-clinical quality profiles were similar among both products, but clinical efficacy and safety must be tested in clinical trials. The technological strategy we describe here can be adapted by other producers, particularly in low- and middle-income countries.

## 1. Introduction

COVID-19 (Coronavirus disease 2019) is a recent pandemic disease caused by the newly emerged severe acute respiratory syndrome coronavirus 2 (SARS-CoV-2)^1^. It has generated millions of infections and hundreds of thousands of deaths (https://covid19.who.int/), and has become a serious threat to global public health and economies worldwide. General symptoms of COVID-19 are fever, severe respiratory illness, dyspnea, and pneumonia, with additional possible complications such as multiple-organ dysfunction or failure, that compromises the gastrointestinal, cardiovascular, renal, and central nervous systems, and can lead to septic shock^2–4^.

Like other coronaviruses, the SARS-CoV-2 virus is an enveloped single-stranded RNA virus, with a virion composed of at least four structural proteins: spike (S), envelope (E), membrane (M), and nucleocapsid (N)^5^. Protein S is implicated in cellular recognition, fusion, and entry^5,6^, making it the most attractive target for SARS-CoV-2 therapeutic development.

Although contagion by this emerging virus continues to increase across the globe, more time is needed to develop and validate vaccines, drugs, and therapies to counteract the disease. Immunoglobulin-based therapies, such as treatment with human convalescent plasma, formulations of immunoglobulins purified from plasma of convalescent patients or hyperimmunized animals, or recombinant monoclonal antibodies, arise as a feasible option that may be achievable on a shorter time scale^7–10^. Treatment of COVID-19 with convalescent plasma seems to be well-tolerated, reduces mortality, and has the potential to improve clinical outcomes, based on several case series studies, matched-control studies, and small randomized clinical trials^11–15^. However, these results must be confirmed by large and ongoing controlled clinical trials.

Formulations of immunoglobulins purified from plasma of convalescent patients have also been used to treat severe respiratory illnesses with viral etiology, but to a lesser extent^7,16,17^. These formulations are advantageous over unpurified convalescent plasma because they are safer and have higher activity, polyvalency, and product consistency^7,9,18^. However, this strategy is donor-dependent, requires strict donor screening for both human pathogens and high levels of neutralizing anti-SARS-CoV-2 antibodies, and relies on well-established blood bank systems that may be scarce in developing countries^8^.

On the other hand, formulations of animal-derived immunoglobulins, such as anti-SARS-CoV F(ab’)2-equine formulations, have shown neutralizing efficacy in cell culture and in *in vivo* murine and hamsters models, as well as in both prophylactic and therapeutic experimental settings^19–23^. Similarly, Zhao et al.^24^ demonstrated effective *in vitro* and *in vivo* neutralization of MERS-CoV by IgG and F(ab’)2-formulations obtained from horses immunized with MERS-CoV virus-like particles (VLPs) expressing MERS-CoV proteins. More recently, two independent groups obtained F(ab’)2 preparations through immunization of horses with recombinant SARS-CoV-2 RBD (receptor binding domain; located at S1 subunit), and demonstrated preclinical efficacy as a potential therapy for COVID-19 both *in vitro*^25,26^ and *in vivo* in a murine model^25^. Also, a F(ab’)2 formulation with high *in vitro* neutralizing potency was developed by immunizing horses with recombinant pre-fusion trimers of SARS-CoV-2 S protein, comprising S1 and S2 subunits^27^.

In light of these favorable outcomes, and harnessing our experience in manufacturing equine snake antivenom^28^, we developed two formulations of whole-IgG from plasma of horses immunized with one of two types of SARS-CoV-2 recombinant proteins: S1 (anti-S1) or a mixture of S1, SEM mosaic, and N proteins (anti-Mix). We detail the manufacturing procedure, quality and safety profiles, and *in vitro* preclinical efficacy of the final formulations with the aim of providing an effective, safe, and affordable potential treatment for COVID-19.

## 2. Methods

### 2.1 Ethical Statement

All procedures involving animals in this study were approved by the Institutional Committee for the Care and Use of Laboratory Animals (CICUA) of the University of Costa Rica (Act 200-2020), and meet both ARRIVE Guidelines^29^ and International Guiding Principles for Biomedical Research Involving Animals^30^. COVID-19 convalescent human plasma collection and use was approved by the Central Committee of Pharmacotherapy (Act GM- CCF-1854-2020) ant the Institutional Bioethics Committee of the Caja Costarricense de Seguro Social (C.C.S.S.; Costa Rican Social Security Fund). All methods were carried out in accordance with relevant regulations stated by the Technical Guidelines for Collection and Convalescent Plasma Processing COVID-19 (PCC), emitted by the C.C.S.S., the Costa Rican Ministry of Health and the University of Costa Rica. All donors were over 18 years old and provided an informed consent approved by the Institutional Committee of Health Record of the C.C.S.S.

### 2.2 Virus proteins

SARS-CoV-2 Spike S1 protein (code REC31828), SARS-CoV-2 Nucleocapsid protein (code REC31812), and SARS-CoV-2 Spike E-M mosaic protein (code REC31829) were purchased from The Native Antigen Company (Oxford, United Kingdom). SARS-CoV-2 Spike RBD Protein (Cat. PNA004) was purchased from Sanyou Biopharmaceuticals Co. Ltd. (Shanghai, China).

### 2.3 Production of anti-SARS-CoV-2 hyperimmune plasma

#### 2.3.1 Hematology, plasma chemistry, and general clinical status of horses

We evaluated the health condition of horses before, during, and after immunization and bleeding. Blood samples were collected from the jugular vein in EDTA-coated vials and electrolytes were measured using an electrolyte analyzer (9180, Roche; Indianapolis, USA). The plasma chemical profile was assessed with a Cobas analyzer (C111, Roche), and hematocrit and hemoglobin concentration values were obtained using a hematology system for veterinary use (EOS, EXIGO; Spånga, Sweden).

#### 2.3.2 Immunization of horses

Two groups of three Ibero-American and mixed breed horses, ranging from 3 to 15 years old and 350 to 450 kg, were immunized with SARS-CoV-2 proteins. The first group (anti-S1) was immunized with the S1 protein alone, while the second group (anti-Mix) was immunized with a mixture of equal parts of S1, N, and Spike-E-M mosaic proteins (SEM). In brief, horses were injected subcutaneously at two weeks intervals according to the scheme summarized in Table 1. Freund's complete adjuvant (F5881, Sigma-Aldrich; Missouri, USA), Freund's incomplete adjuvant (F5506-Sigma-Aldrich), and Emulsigen-D adjuvant (MVP adjuvants; Nebraska, USA) were included to enhance the antibody response of horses. Samples of serum were collected before and during immunization with each booster and stored at −20 °C until use. Antibody responses of horses to SARS-CoV-2 proteins was monitored via ELISA as described below.

**Table 1.**
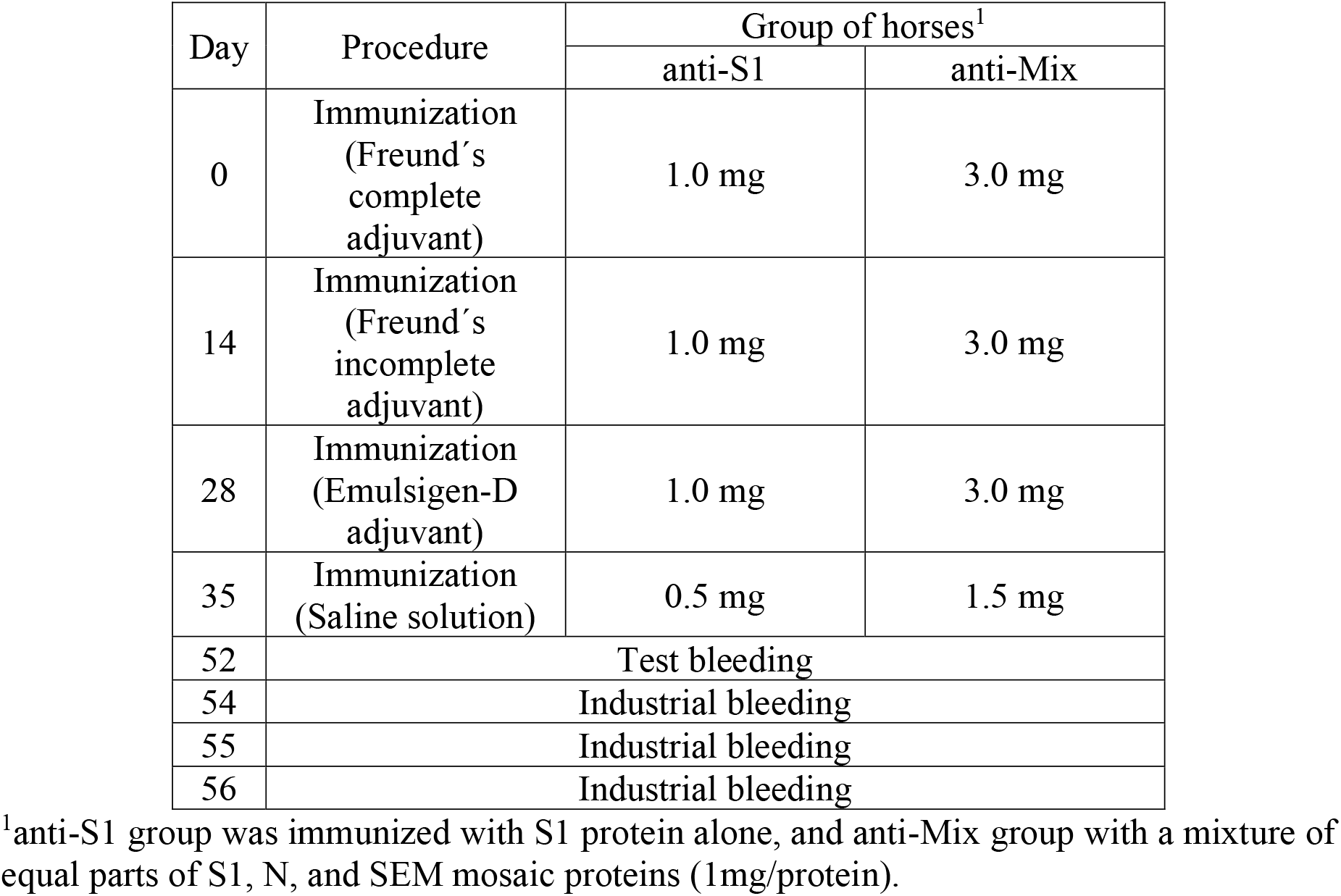
Immunization and bleeding protocol for anti-SARS-CoV-2 hyperimmune plasma production.

#### 2.3.3 Bleeding of horses and plasma separation

We bled horses 19 days after the last injection of viral proteins using a closed system of blood collection bags. In total, three blood collections of 6 L each were performed across three consecutive days. The first day, blood was collected and stored overnight at 2-8 °C to allow erythrocyte sedimentation. The second day, plasma from the first day was separated from erythrocytes, preserved by the addition of 0.005 % thimerosal, and stored at 2-8 °C until use. Erythrocytes were suspended in saline solution and warmed to 37 °C. Then, following the second collection of blood, tempered erythrocytes from the first day were transfused back to the same animal. The third day, we repeated the same procedure as on the second day^31^.

### 2.4 Purification and formulation of immunoglobulins

To prepare anti-S1 and anti-Mix formulations, we purified immunoglobulins using the caprylic acid precipitation method^32^. Immunoglobulins were formulated at 65 g/L total protein, 8.5 g/L NaCl, 2.0 g/L phenol and pH 7.0. Then, purified antibodies were sterilized by 0.22 μm pore membrane filtration and dispensed in 10 mL glass vials.

### 2.5 Physicochemical analysis

#### 2.5.1 Electrophoretic analysis

Electrophoretic analyses of viral proteins and equine formulations were performed in 4-20% SDS-PAGE gel Mini-PROTEAN TGX (BioRad; California, USA) and 7.5% SDS-polyacrylamide gels (SDS-PAGE), respectively. All samples were analyzed under nonreducing conditions^33^, and compared with Thermo Scientific (26616; Massachusetts, USA) pre-stained molecular mass markers. Protein bands were stained with Coomassie Brilliant Blue R-250, and gels were destained with a mixture of methanol, ethanol, and acetic acid.

#### 2.5.2 Total protein concentration

Total protein concentration was determined using a modification of the Biuret test^34^. Fifty microliters of protein standards were mixed with 2.5 mL of Biuret reagent and incubated at room temperature for 30 min. We created a calibration curve with absorbances of the standards at 540 nm. Then, we repeated the procedure with samples and calculated protein concentration based on the equation of the calibration curve.

#### 2.5.3 IgG monomer content

The content of IgG monomers was assessed by FPLC gel filtration chromatography in a Superdex 200 10/300 GL column (GE Healthcare, Pharmacia; Stockholm, Sweden), using 0.15 mol/L NaCl, 20 mmol/L Tris, pH 7.5 as the mobile phase with a 0.5 mL/min flow. Protein peaks were detected by measuring absorbance at 280 nm.

#### 2.5.4 Turbidity

Turbidity was quantified using a turbidimeter (220, La Motte; Maryland, USA) calibrated with HACH Stablcal® turbidity standards prior to analysis. Turbidity was expressed in nephelometric turbidity units (NTU).

#### 2.5.5 pH

We measured pH with a pH meter (Orion 4 Star, ThermoScientific; Massachusetts, USA) equipped with a glass electrode.

#### 2.5.6 Sodium chloride content

Sodium chloride was quantified according to the Pharmacopeial methodology^35^. In brief, 5.0 mL of each sample was diluted with distilled water, mixed with acetic acid and methanol, and titrated with a standard solution of 0.1 mol/L silver nitrate. Eosin Y was used as the indicator.

#### 2.5.7 Osmolality

Osmolality was assessed by a cryoscopic technique using a micro-osmometer (3320, Advanced Instrument; Massachusetts, USA) and a reference solution.

#### 2.5.8 Phenol concentration

Phenol concentration was determined by a colorimetric assay^36^, using 4-aminoantipyrine and potassium ferricyanide to form a derivative compound that absorbs at 505 nm.

#### 2.5.9 Caprylic acid concentration

Caprylic acid concentration was quantified by RP-HPLC following Herrera et al.^37^ using a liquid chromatographer (1220, Agilent Technologies; California, USA) equipped with an Eclipse XDB-C8 5 μm column (150 mm × 4.6 mm i.d.). The mobile phase was a mixture of acetonitrile and water (60/40, v/v) and the flow rate was 1 mL/min. The UV detection was set at 210 nm.

### 2.6 Microbiological analysis

#### 2.6.1 Endotoxin assay

Endotoxins were detected and quantified using the gel clot method as described by Solano et al.^38^. We added 0.2 mL of in-process and final product sample dilutions to single test vials of *Limulus* amoebocyte lysate (LAL; Cat # 65003 Pyrotell®, ACC; Massachusetts, USA). Dilutions of lipopolysaccharide standard (LPS; Cat # E0005, ACC) prepared with samples were used as positive controls, and LAL Reagent Water (Cat # WP1001, ACC) was used as a negative control. After 1h incubation at 37 °C, tubes were gently inverted 180° to assess gelation of the mixture.

#### 2.6.2 Sterility test

Sterility test was conducted according to USP 37^35^. A predetermined volume of anti-S1 and anti-Mix products was aseptically filtered through a 0.22 μm pore membrane. Membranes were cut into equal sizes, and each half was transferred to one of two types of culture media suitable for the growth of fungi as well as aerobic and anaerobic microorganisms. After inoculation, media were incubated for 14 days at either 25 °C or 35 °C depending on the media type. During and at the end of the incubation period, media were examined for macroscopic evidence of microbial growth. Sterility compliance was dependent on the absence of microbial growth.

### 2.7 Immunological analysis

#### 2.7.1 Human COVID-19 convalescent plasma

Human convalescent plasma was used as a control. We used a mixture of plasma voluntarily donated by 28 recovered SARS-CoV-2 patients. All patients provided informed consent and tested positive for anti-SARS-CoV-2 antibodies with (1) anti-SARS-CoV-2 ELISA IgG kit (EI 2606-9601 G, EUROIMMUN; Lübeck, Germany), (2) MAGLUMI 2019-nCoV (SARS-CoV-2) IgM and IgG kits (SNB-130219015M and SNB-130219016M, Snibe Diagnostics; Shenzhen, China), and (3) Standard Q COVID-19 IgM/IgG Combo Test Kit (09COV50G, SD Biosensor; Gyeonggi, Korea). All tests were performed according to manufacturer’s instructions. The pool of convalescent plasma was freeze-dried and stored at −20 °C until use.

#### 2.7.2 ELISA

Polystyrene plates (Costar 9017, Corning Inc.; New York, USA) were coated overnight at room temperature with 0.5 μg/well of S1 protein, 0.25 μg/well of nucleocapsid protein, 1 μg/well of mosaic protein or 0.5 μg/well of RBD protein, depending on the experiment and in duplicate. After washing the plates five times with distilled water, 100 μL of equine plasma or immunoglobulin formulation, diluted with 2% skim milk/PBS, were added to each well. The plates were incubated for 1 h at room temperature and washed again five times. Afterwards, 100 μL of rabbit anti-equine IgG antibodies conjugated with peroxidase (A6917, Sigma-Aldrich), diluted 1:3000 or 1:5000 with 2% skim milk/PBS, were added to each well. Again, microplates were incubated for 1 h at 25 °C. After a final washing step, color was developed by the addition of H2O2 and *o*-phenylenediamine as a substrate (P9029, Sigma-Aldrich). Color development was stopped by the addition of 2.0 mol/L HCl. Absorbances were recorded at 492 nm. For ELISA titration of final formulations, titer was calculated as the dilution at which the absorbance was equal to five times the absorbance of a purified normal equine plasma (normal equine immunoglobulins, NEI) diluted 1:1000.

#### 2.7.3 Western blot

For Western blot analysis, viral proteins were separated by SDS-PAGE as described above. Then, proteins were transferred to a nitrocellulose membrane and blocked with 1% skim milk/PBS for 40 min at room temperature. The membrane was incubated with a dilution 1:1000 of either anti-S1 or anti-Mix samples in 0.1% skim milk/PBS for 1 h at 25 °C. Subsequently, a second incubation was performed with a dilution 1:1000 of a rabbit anti-equine IgG antibody conjugated with peroxidase. A precipitating chromogenic substrate (4- Chloro-1-Naphthol; C6788, Sigma-Aldrich) was added.

#### 2.7.4 Plaque Reduction Neutralization (PRNT)

Plaque Reduction Neutralization for anti-S1 and anti-Mix formulations and human convalescent plasma was determined by assessing the ability of the samples to neutralize SARS-CoV-2 virus (2019-nCoV/USA-WA1/2020). PRNT was performed at BSL-3 facilities. Samples were heat-inactivated for 30 minutes at 56 °C, and then diluted to appropriate concentrations in Dulbecco's Modified Eagle's Medium (DMEM) (VWRL0102- 500, VWR Life Sciences; Pennsylvania, USA), supplemented with 5 % heat-inactivated fetal bovine essence (FBE) (VWR10803-034, VWR Life Sciences), 1 % Penicillin/Streptomycin (P/S) (15-140-122, Gibco Thermo Scientific) and 1 % L-Glutamine (VWRL0131-0100, VWR Life Sciences). SARS-CoV-2 was diluted in supplemented DMEM to appropriate concentration. Virus was then added to antibody samples and allowed to incubate for 1 hour at 37 °C and 5 % CO2. After incubation, viral plaque assay was conducted to quantify viral titers. 12-well plates were previously seeded with Vero cells (ATCC CCL-81; Virginia, USA) at a density of 2 x 10^5^ cells per well. Plates were inoculated for 1 hour at 37 °C and 5 % CO2. After infection, a 1:1 overlay consisting of 0.6 % agarose and 2X Eagle’s Minimum Essential Medium without phenol red (115-073-101, Quality Biological; Maryland, USA), supplemented with 10 % fetal bovine serum (FBS) (10-437-028, Gibco Thermo Scientific), non-essential amino acids (11140-050, Gibco Thermo Scientific), 1 mM sodium pyruvate (25-000-Cl, Corning Inc.), 2 mM L-glutamine, 1 % P/S was added to each well. Plates were incubated at 37 °C for 48 hours. Cells were fixed with 10 % formaldehyde for 1 hour at room temperature. Formaldehyde was aspirated and the agarose overlay was removed. Cells were stained with crystal violet (1 % w/v in a 20 % ethanol solution). Viral titer of SARS-CoV-2 was determined by counting the number of plaques. Each sample was analyzed in triplicate. The median effective dose (ED50) was calculated using Probit analysis and expressed as the dilution factor of the formulations in which 50 % of the virus was detected. Alternatively, neutralization was expressed as the ratio of protein concentration/ED50 (μg/mL).

#### 2.7.5 IgG-mediated activation of human Fcγ receptors (FcγR)

Activation of human FcγRIIIA (CD16) was assessed using a surrogate activation assay^39^. Briefly, IgG-dependent activation of BW:FcγRIIIA-ζ transfectants, i.e. BW5147 thymoma cells (TIB-47TM ATCC; Virginia, USA), expressing the extracellular portion of human FcγRIIIA (higher affinity variant with valine in position 158) fused to the mouse CD3 ζ-chain, was measured. SARS-CoV-2 S1 and N proteins were coated on plates using coating buffer (0.1 mol/L Na2HPO4, pH 9.0). After blocking with 5 % fetal calf serum (FCS), 20 mg/dL of either anti-S1, anti-Mix, or convalescent human plasma in DMEM 10 % (v/v) FCS was added for 30 min at 37 °C in an atmosphere of 5% CO2. To remove non-immune IgG, plates were washed three times with DMEM containing 10% (v/v) FCS. Then, 100 000 BW:FcγRIIIA-ζ reporter cells per well in RPMI 10 % (v/v) FCS medium were added. After co-cultivation for 16 h at 37 °C in a 5 % CO2 atmosphere, supernatants were diluted 1:2 in ELISA sample buffer (PBS with 10 % [v/v] FCS and 0.1 % [v/v] Tween-20) and mIL-2 was measured by ELISA using the capture Ab JES6-1A12 and the biotinylated detection Ab JES6-5H4 at 450 nm (BD Pharmingen™; San Diego, USA). Experiments were performed in triplicate.

### 2.8 Statistical Analysis

Statistical analyses were performed using the software IBM® SPSS® Statistics v24 (New York, USA). Results were expressed as means ± SD. A paired t-test was used to evaluate hematocrit and hemoglobin values of horses before and after immunization. A repeated measures ANOVA was used to investigate changes in the biochemical parameters of the horses during plasma production, and values were corrected by the Greenhouse-Geisser factor when needed. A two-way independent ANOVA was conducted to assess ELISA absorbance of formulations and antigens. A Bonferroni post-hoc analysis was performed to elucidate differences among groups. Additionally, a bootstrap BCa 95% CI was performed to validate the assumption of the equality of variances. All *P*-values < 0.05 were considered significantly different. ED50 values were considered significantly different if confidence intervals at 95% did not overlap.

## 3. Results and discussion

### 3.1 SARS-CoV-2 recombinant proteins

Structural SARS-CoV-2 recombinant S1, N, and SEM mosaic proteins were used as immunogens for the preparation of equine formulations. S1 forms part of the transmembrane spike (S) glycoprotein homotrimer located at the viral surface. S is composed of S1 and S2 subunit domains, and is the main protein responsible for recognition of the angiotensin-converting enzyme 2 (ACE2) receptor via the receptor binding domain (RBD) at the N-terminal S1 subunit^5,6^. Thus, S is the most suitable target for immunoglobulin-based therapy for COVID-19^1^,^40^. Both S1 and S2 subunits of SARS-CoV, MERS-CoV, and SARS-CoV-2, specially conformational epitopes at S1, and particularly at RBD^41–44^, have served as targets for developing neutralizing antibodies to disable receptor interactions and block viral entry to host cells. Also, during SARS-CoV infection, it was demonstrated that protective humoral and T-cellular immunity is induced by S protein^45^.

Nucleocapsid phosphoprotein (N) plays an essential role in genome packaging, virion assembly, replication, and transcription^46^. This protein is highly expressed during infection^47^, and several serological studies of COVID-19 convalescent patients have reported seropositivity against N^47–51^. Antibody response to N protein is similar to RBD reactivity^52^, and therefore may contribute to some extent of the preliminary efficacy seen with human convalescent plasma therapies^15^.

Membrane (M) protein is the most abundant of the structural proteins in the virus. This transmembrane protein coordinates virus assembly by interacting with the other structural proteins^53^. In contrast, the role of the envelope (E) protein is not entirely understood; this viroporin is expressed profusely during viral replication cycles, and has been implicated in viral assembly, budding, envelope formation, and pathogenesis^53,54^. Immunization of horses with VLPs expressing MERS-CoV S, M and E proteins has successfully induced neutralizing antibodies against MERS-CoV^24^.

Here, we used SARS-CoV-2 recombinant S1 protein (produced in baculovirus insect cells), N protein (expressed in *E. colĩ),* and SEM mosaic (an *E. coli* derived recombinant protein containing the S, E, and M immunodominant regions) as immunogens to produce two formulations of equine immunoglobulins: anti-S1 (towards S1 protein) and anti-Mix (towards a mixture of S1, N, and SEM mosaic proteins). Additionally, we used recombinant RBD (expressed in HEK293 cells) for immunoreactivity and quality control assessment.

Before immunization, we verified the purity of viral protein preparations with SDS-PAGE analysis (Fig. 1) and found that S1 protein was represented by a ~85 kDa band of high purity (94%). N protein showed a band of ~57 kDa and 75% purity. SEM mosaic protein presented only two bands at 85 kDa (S protein) and 13 kDa (E protein). RBD protein showed a 34 kDa band that represented 95% of the preparation. The identity of all recombinant proteins was confirmed by mass spectrometry (nESI-MS/MS; results not shown). Toxicity of recombinant proteins used as immunogens was assessed in a murine model, where no evidence of toxicity was observed (results not shown).

**Figure 1.**
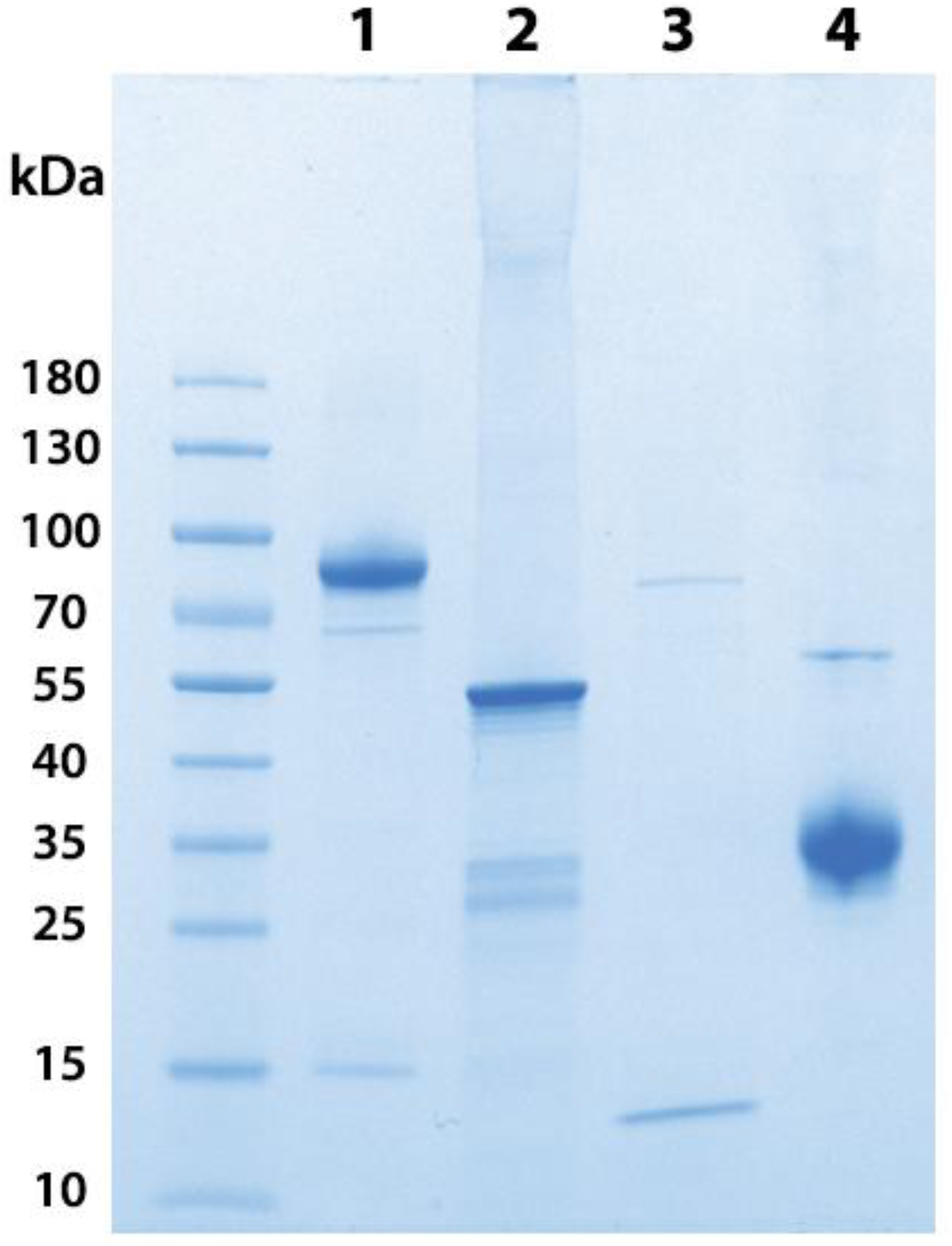
SDS-PAGE analysis of SARS-CoV-2 recombinant proteins. Lane 1: S1; Lane 2: N; Lane 3: SEM mosaic; Lane 4: RBD. Samples (5 μg) were loaded in a 4-20% polyacrylamide gradient gel in the presence of SDS and run under non-reducing conditions. The gel was stained with Coomassie Blue.

### 3.2 Production of anti-SARS-CoV-2 hyperimmune plasma

#### 3.2.1 Immunization, antibody response, and clinical evaluation of horses

An ideal design for an adequate immunization strategy takes into consideration several factors. For example, the immunogenicity of proteins is closely related to their structural characteristics such as molecular mass, tertiary and quaternary structure, and post-translational modifications. Similarly, optimal immunization procedures can vary depending on the animal model, administration route, use of immunological adjuvants, dose, and frequency of boosters. After the first injection of foreign proteins, naïve lymphocytes are recruited and activated by antigen-presenting cells, leading to the proliferation and creation of memory T and B cells. In turn, plasma B cells produce specific antibodies, whose affinity and specificity most likely mature after each booster^55^. In this experiment, immunization of horses followed the scheme summarized in Table 1. During immunization, antibody responses varied among individual horses. In general, antibody titers started to increase 15 days after priming, and reached a plateau at the maximum anti-viral antibody level around 7 weeks (52 days; Fig. 2). Others have reported maximum antibody responses 6-7 weeks after immunizing horses with SARS-CoV particles^20,21^, and 4-6 weeks after immunizing horses with SARS-CoV-2 RBD^25,26^ and trimeric S protein^27^. In this study, horses in anti-S1 and anti-Mix groups showed similar dynamics in their development of plasma concentrations of antibodies towards S1 (Fig. 2a), which is not surprising because both groups received the S1 immunogen at the same dosage, schedule, and method of administration. Antibodies towards N protein were only developed by horses in the anti-Mix group (Fig. 2b).

**Figure 2.**
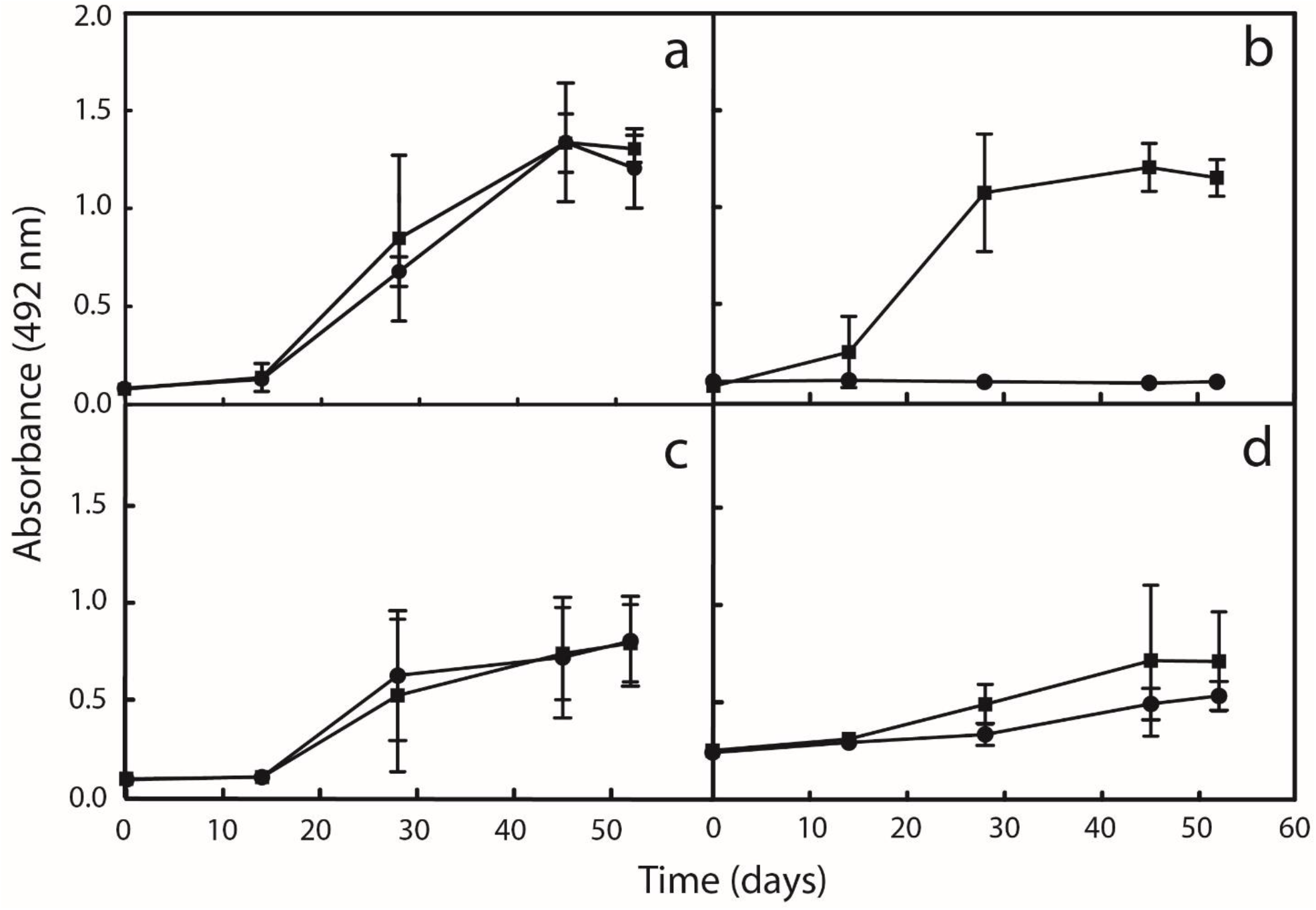
ELISA antibody response of pools of plasma from the anti-S1 and anti-Mix groups of horses during the immunization schedule with SARS-CoV-2 recombinant proteins. a: anti-S1 response; b: anti-N response; c: anti-SEM mosaic response; d: anti-RBD response. Anti-S1 group is represented by a circle (•) and anti-Mix group is represented by a square (■). Samples towards S1 and N proteins were assessed at a 1/1000 dilution, and samples towards SEM mosaic and RBD were assessed at a 1/500 dilution. Absorbances were recorded at 492 nm. Results are expressed as mean ± SD.

Because RBD is a domain of S1^5,6^, we anticipated and observed that both groups would show similar responses towards RBD, even though it was not used as an immunogen (Fig. 2c). Also, despite the fact that only the anti-Mix group was immunized against the SEM mosaic, both groups of horses developed a similar antibody response towards this immunogen (Fig. 2d); presumably, the S immunodominant region contained in the mosaic masked the responses to E and M proteins.

Throughout the immunization schedule, animals used for hyperimmune plasma production should be under rigorous veterinary care, ensuring that the health and welfare of each animal are closely monitored, and ethical guidelines are appropriately met. During this process, the stimulation of a local inflammation is expected for the production of a cellular infiltrate, which at the beginning is mainly composed of neutrophils and at the end by antigen-presenting cells^55^.

In our study, horses developed local inflammation and signs of minor pain and discomfort such as lameness or abnormal gait, which lasted in most cases two days after each booster. However, the use of an anti-inflammatory drug was necessary for pain relief in two horses in the anti-Mix group. The complete and incomplete Freund’s adjuvants produced fistulation with minor pus-like discharge between 2 and 3 weeks after each injection. In spite of their proven adjuvant efficacy and depot effect, it is recommended to limit the use of these adjuvants to the beginning of the immunization protocol because they are known to cause local injuries^56^. Therefore, for the third booster, we selected Emulsigen-D adjuvant, which only produced some ventral edema on anti-Mix horses.

At the end of the immunization scheme, the immunogen dose was reduced from 1 mg to 0.5 mg of each immunogen to reduce inflammatory effects and to ration immunogen use. Overall, milder inflammation was observed in the anti-S group than in the anti-Mix group. This difference may result from protein interactions in the anti-Mix preparation and/or the fact that the anti-Mix group received 3 mg of total immunogen versus only 1 mg in the anti-S1 group. Nonetheless, further studies are necessary to confirm these hypotheses.

Hematological parameters were analyzed regularly throughout the immunization scheme, and before bleeding (Table 2). Slight decreases in hematocrit and hemoglobin were observed in some horses, but were not statistically significant (F= 0.550, df= 5, P= 0.606; F= 1.179, df= 5, P= 0.291, respectively). Nonetheless, the veterinary team treated horses with an antianemic vitamin complex (Complemil500®, Kyrovet Laboratories; Bogotá, Colombia) to ensure a safe blood extraction process.

**Table 2.** Hematological and serum biochemical parameters of anti-S1 and anti-Mix groups of horses during immunization with SARS-CoV-2 recombinant proteins^1^.

| Parameter | Anti-S1 group |  |  | Anti-Mix group |  |  |
| --- | --- | --- | --- | --- | --- | --- |
|  | Pre-immunization | Before bleeding | After bleeding | Pre-immunization | Before bleeding | After bleeding |
| Hematocrit (%) | 38.7 ± 8.1 | 34.0 ± 4.2 | ND<sup>2</sup> | 34.4 ± 5.6 | 35.4 ± 10.9 | ND<sup>2</sup> |
| Hemoglobin (g/dL) | 13.4 ± 2.7 | 11.3 ± 1.3 | ND<sup>2</sup> | 12.1 ± 1.4 | 11.8 ± 3.0 | ND<sup>2</sup> |
| Total protein (g/L)* | 57.0 ± 2.6 | 68.7 ± 9.5 | 61.3 ± 2.3 | 58.7 ± 7.0 | 74.0 ± 10.0 | 65.3 ± 3.1 |
| Albumin (g/L)* | 24.6 ± 0.9 | 20.3 ± 4.6 | 22.2 ± 1.6 | 22.6 ± 0.8 | 20.6 ± 1.7 | 22.7 ± 1.9 |
| Globulins (g/L)* | 32.3 ± 2.9 | 48.4 ± 5.2 | 39.2 ± 4.0 | 36.1 ± 7.8 | 53.4 ± 10.4 | 42.6 ± 4.3 |
| Sodium (mmol/L) | 135.0 ± 1.7 | 133.3 ± 15.3 | 137.0 ± 0.0 | 133.0 ± 1.7 | 135.3 ± 7.2 | 136.7 ± 0.6 |
| Potassium (mmol/L)* | 4.2 ± 0.2 | 2.7 ± 0.2 | 3.7 ± 0.2 | 4.1 ± 0.1 | 2.9 ± 0.6 | 4.3 ± 0.1 |
| Chloride (mmol/L)<sup>3</sup> | 101.0 ± 1.7 | 96.3 ± 12.9 | 106.0 ± 1.0 | 97.7 ± 0.6 | 96.3 ± 4.6 | 104.0 ± 1.0 |
| Phosphorus (mmol/L)* | 0.7 ± 0.1 | 1.2 ± 0.1 | 0.9 ± 0.4 | 0.8 ± 0.1 | 1.5 ± 0.6 | 1.1 ± 0.3 |
| Calcium (mmol/L)* | 2.8 ± 0.3 | 2.3 ± 0.1 | 2.2 ± 0.1 | 2.5 ± 0.2 | 2.4 ± 0.2 | 2.3 ± 0.1 |
| Bicarbonate (mmol/L)<sup>3</sup> | 21.3 ± 2.1 | 20.6 ± 2.1 | 14.0 ± 12.0 | 21.7 ± 1.7 | 18.8 ± 2.6 | 20.4 ± 0.9 |
| Creatinine (μmol/L) | 85.4 ± 9.9 | 76.8 ± 12.8 | 79.8 ± 1.9 | 68.3 ± 10.3 | 73.9 ± 14.0 | 72.7 ± 8.4 |
| Urea (mmol/L) | 5.9 ± 0.8 | 6.1 ± 1.1 | 5.7 ± 0.7 | 5.2 ± 0.7 | 6.2 ± 1.2 | 6.4 ± 0.7 |
| GGT (U/L) | 14.9 ± 3.7 | 21.1 ± 10.6 | 11.1 ± 8.3 | 19.8 ± 21.7 | 38.9 ± 44.8 | 33.4 ± 37.3 |
| AST (U/L) | 359.7 ± 13.2 | 350.0 ± 90.5 | 383.4 ± 113.0 | 290.6 ± 28.6 | 330.6 ± 94.5 | 288.2 ± 58.1 |
| CK (U/L) | 221.8 ± 20.5 | 470.4 ± 272.4 | 527.8 ± 210.2 | 345.8 ± 114.1 | 484.8 ± 232.3 | 391.0 ± 24.8 |
| Glucose (mmol/L)<sup>3</sup> | 5.4 ± 0.8 | 3.5 ± 0.9 | 6.1 ± 2.6 | 5.6 ± 0.5 | 3.1 ± 0.3 | 6.7 ± 3.6 |

<sup>1</sup> Results are expressed as mean ± SD (n = 3 horses).
<sup>2</sup> Not determined.
<sup>3</sup> Did not comply with the assumption of sphericity; corrected with Greenhouse-Geisser factor.
GGT: Gamma-Glutamyl Transferase; AST: Aspartate Aminotransferase; CK: Creatine Kinase.
\*Parameter changed significantly different over time (P < 0.05).

In general, biochemical analysis of horse serum showed normal values, although some measures varied significantly during hyperimmune plasma production (Table 2; P < 0.05). Pre-bleeding analysis revealed a significant increase in total protein and globulins and a decrease of albumin as a direct consequence of hyperimmunization and production of anti-SARS-CoV-2 antibodies. Potassium values decreased below the normal range, an observation also reported by Angulo et al.^31^ in horses injected with snake venoms for antivenom manufacture. Although there was a slight rise in the levels of gamma glutamyl transferase (GGT) and creatine kinase (CK) enzymes, which may reflect some hepatic and tissue damage. Nonetheless, those changes were not significantly different than baseline serum values (P > 0.05). Results suggest a possible effect of virus proteins or adjuvants on the electrolyte profile and protein balance of plasma producing animals.

#### 3.2.2 Bleeding for hyperimmune plasma collection

We bled horses when the maximum anti-viral antibody level against all immunogens reached a plateau (Fig. 2). First, horses underwent physical examination, including evaluation of body condition, auscultation of cardiorespiratory and digestive systems, and hematological tests. Strict veterinary surveillance was maintained throughout bleeding, self-transfusion, and post-bleeding.

Bleeding was performed across three consecutive days. Horses had access to water and food *ad libitum.* By the end of the process, an average of 9.2 ± 1.0 L plasma was collected from each horse. Bleeding and plasma separation were performed by specialized technicians. Laboratories, equipment, and a closed system of blood collection bags were designed to operate with refined aseptic technique. All the plasma bags tested negatively to endotoxins, containing less than 1.5 EU/mL, and met all the specifications required to be included in the plasma pools for the immunoglobulin purification process.

Bleeding resulted in no adverse acute or chronic physiological alterations. Some minor shifts were observed in serum biochemical parameters, but within accepted ranges (Table 2) as reported by Cunha et al.^27^. Horses showed an expected decrease in total protein and globulins concentration after bleeding, as a consequence of the removal of plasma proteins. After a 2-month rest period, animals recovered to normal status, and were ready to initiate a new cycle of immunization and bleeding.

### 3.3 Pilot scale antisera production and quality control of final products

#### 3.3.1 Anti-SARS-CoV-2 immunoglobulin purification

To prepare antisera, hyperimmune plasma was pooled by immunization group (28.9 kg anti-S1 plasma and 26.5 kg anti-Mix plasma), and subsequently fractionated by caprylic acid precipitation. This method is routinely used at the Instituto Clodomiro Picado of the University of Costa Rica for the manufacture of whole-IgG antivenoms, and produces satisfactory yield and adequate purity of preparations^32^. The antivenoms generated using this fractionation protocol have proven safe and effective in clinical trials in patients suffering snakebite envenomings^57–59^.

Caprylic acid has the advantage of precipitating non-IgG proteins from plasma, while leaving the pharmacologically active ingredient - the immunoglobulins - in solution^32^. The advantage of this method is that IgG aggregation is circumvented. After precipitation, insoluble material was removed by filtration, producing a clarified solution enriched in plasma immunoglobulins.

Once purified, whole-IgG preparations were properly formulated in compliance with quality control specifications (Table 3). Solutions were dialyzed and concentrated to remove caprylic acid (Table 3; ≤ 250 mg/L), and to reach final protein concentration. Additionally, pH, osmolality, and ionic strength were adjusted to values compatible with parenteral administration, and phenol was added as a preservative (Table 3).

**Table 3.** Quality control assessment of anti-S1 and anti-Mix immunoglobulin preparations^1^.

| Parameter | anti-S1 | anti-Mix | Specification |
| --- | --- | --- | --- |
| Physicochemical purity (%) | 90.2 | 91.3 | > 90 |
| Total protein (g/dL) | 6.62 ± 0.03 | 6.44 ± 0.05 | ≥ 5 |
| Monomers (%)<sup>2</sup> | 92.57 | 92.57 | > 90 |
| Turbidity (NTU)<sup>3</sup> | 29.36 ± 0.75 | 28.10 ± 0.64 | ≤ 50 |
| pH | 7.03 ± 0.03 | 7.08 ± 0.01 | 6.5-7.5 |
| Chloride (g/dL) | 0.893 ± 0.004 | 0.815 ± 0.006 | 0.75 – 1.0 |
| Osmolality (mOsmol/kg) | 335.7 ± 1.5 | 304.7 ± 1.5 | ≥ 240 |
| Phenol (g/dL) | 0.21 ± 0.00 | 0.204 ± 0.002 | 0.10 - 0.25 |
| Caprylic acid (mg/L) | 218.3 ± 5.0 | 114.4 ± 5.3 | ≤ 250 |
| LAL (EU/mL) | < 3 | < 3 | < 35<sup>4</sup> |
| Sterility | Absence of growth | Absence of growth | Absence of growth |
| Anti-S1 ELISA titer<sup>5</sup> | 19062 ± 139 | 11172 ± 386 | Pending<sup>6</sup> |
| Anti-N ELISA titer<sup>5</sup> | 1:2 ± 1 | 3147 ± 116 | Pending<sup>6</sup> |

<sup>1</sup>Formulations were adjusted to a similar protein concentration.
<sup>2</sup>Monomer content is expressed as the relative percent of monomeric immunoglobulin proteins as analyzed by size exclusion chromatography.
<sup>3</sup>Turbidity is expressed in nephelometric turbidity units (NTU).
<sup>4</sup>Endotoxin limit was calculated as the quotient K/M, where K is the threshold pyrogenic dose of endotoxin per kilogram of body weight (5 EU/kg), and M is the maximum total dose administered to a 70 kg-patient for 1 h. Based on our results, the recommended dose for anti-S1 and anti-Mix formulations is 10 mL. Therefore, the acceptable endotoxin limit for this formulation is: (5 EU/kg)/(10 mL/70 kg) = 35 EU/mL.
<sup>5</sup>ELISA titer was calculated as the dilution at which the sample absorbance was equal to five times the absorbance of a normal equine immunoglobulin preparation (NEI) diluted 1:1000.
<sup>6</sup>This specification will be established after evaluation of the formulations in a clinical study. Data are expressed as mean ± SD.

Finally, a 0.22 μm-filtration was performed prior to aseptic filling of vials as a final sterilization step to avoid contamination with potential pathogens such as bacteria, protozoa, and fungi. After downstream processing of the formulations, we filled 612 and 479 vials containing 10 mL of purified and concentrated anti-S1 and anti-Mix products, respectively. In other words, we obtained approximately 20 vials/L plasma.

Because heterologous formulations are animal plasma-derived products, there is a theoretical concern regarding transmission of viral infectious agents. To prevent this possibility, viral risk assessment needs to be performed via rigorous control of the viral load of the raw material, as well as evaluation of both existing and intentionally introduced antiviral steps during production^56^. The use of caprylic acid in the production of snake antivenoms also works as an antiviral step for enveloped viruses, reducing viral loads by up to 5 log10 for several model viruses^60^. Previously, we have found that precipitation of equine plasma with 6 % caprylic acid reduces infectivity of human herpesvirus (HSV-1) in Vero cells by more than 5 log10 within 5 minutes of adding the precipitating agent (our unpublished data). Moreover, 0.25 % phenol has been reported as an efficient virucidal agent of enveloped viruses when added to final formulations of snake antivenoms^61^. Therefore, our anti-SARS-CoV-2 formulations were produced with a methodology that has been previously validated as having two viral inactivation steps.

#### 3.3.2 Physicochemical and microbiological assessment of final products

A comparison between various physicochemical parameters of both formulations is presented in Table 3. Both preparations complied with quality control specifications. The immunoglobulin monomer content of both products indicated that they have few soluble protein aggregates, and the turbidity due to insoluble aggregates was low. This may be relevant in terms of the safety profile of the product, because the presence of aggregates of immunoglobulins in heterologous formulations has been proposed as one of the main causes of early adverse reactions^62^. Endotoxin and sterility tests were fully compliant, which also supports the microbiological safety profile of the preparations.

Figure 3 (lanes 2 and 4) shows the electrophoretic profiles of the formulations. Both compare favorably in terms of purity (Table 3; > 90%) and presented a predominant band with a molecular mass corresponding to whole IgG ~150 kDa. As a result of the efficacy of purification by caprylic acid, there are only traces of protein contaminants.

**Figure 3:**
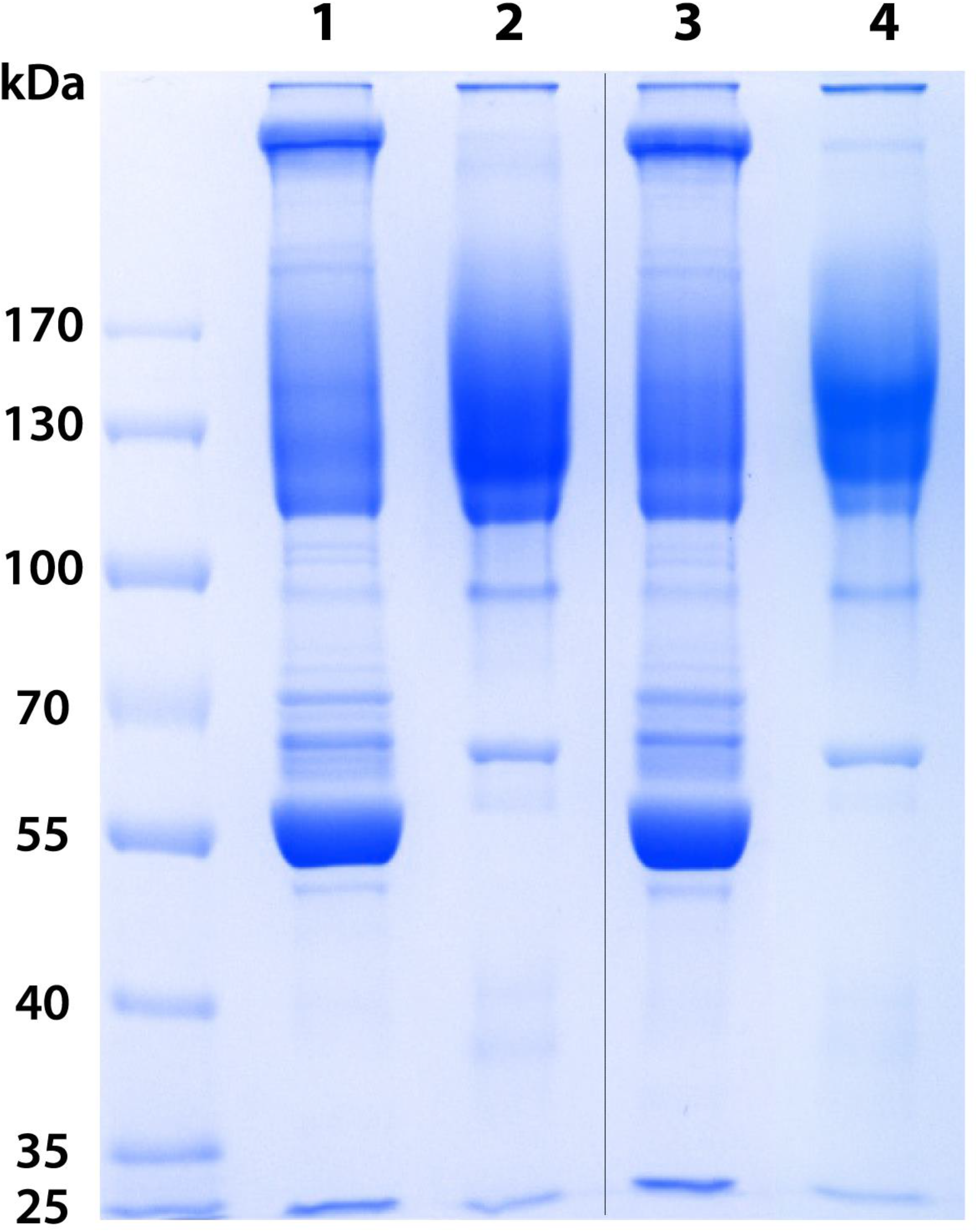
SDS-PAGE analysis of pools of plasma and final anti-S1 and anti-Mix formulations. Lane 1: Hyperimmune pool of plasma anti-S1; Lane 2: Anti-S1 formulation; Lane 3: Hyperimmune pool of plasma anti-Mix; Lane 4: Anti-Mix formulation. Samples (20 μg) were loaded in a 7.5% polyacrylamide gel in the presence of SDS and run under nonreducing conditions. The gel was stained with Coomassie Blue. The original gel was cropped; the full-length gel is presented in Supplementary Figure S1.

#### 3.3.3 Immunological assessment of final products

##### 3.3.3.1 Immunoreactivity

The ELISA antibody response of anti-S1 and anti-Mix formulations towards the four recombinant proteins used as immunogens was significantly different (F= 797.529, df= 2;24, P< 0.0001; Fig. 4a). Such immunoreactivity agrees with that seen during horse immunization, as the anti-S1 formulation recognized both S1 and RBD and the anti-Mix formulation recognized S1, N, and RBD. Anti-RBD immunoreactivity was higher with the anti-Mix formulation (P < 0.05), whereas the SEM mosaic was poorly recognized by both anti-S1 (x = 0.118 [0.101 − 0.146] 95% Bca) and anti-Mix (x= 0.198 [0.191 - 0.203] 95% Bca) formulations.

**Figure 4.**
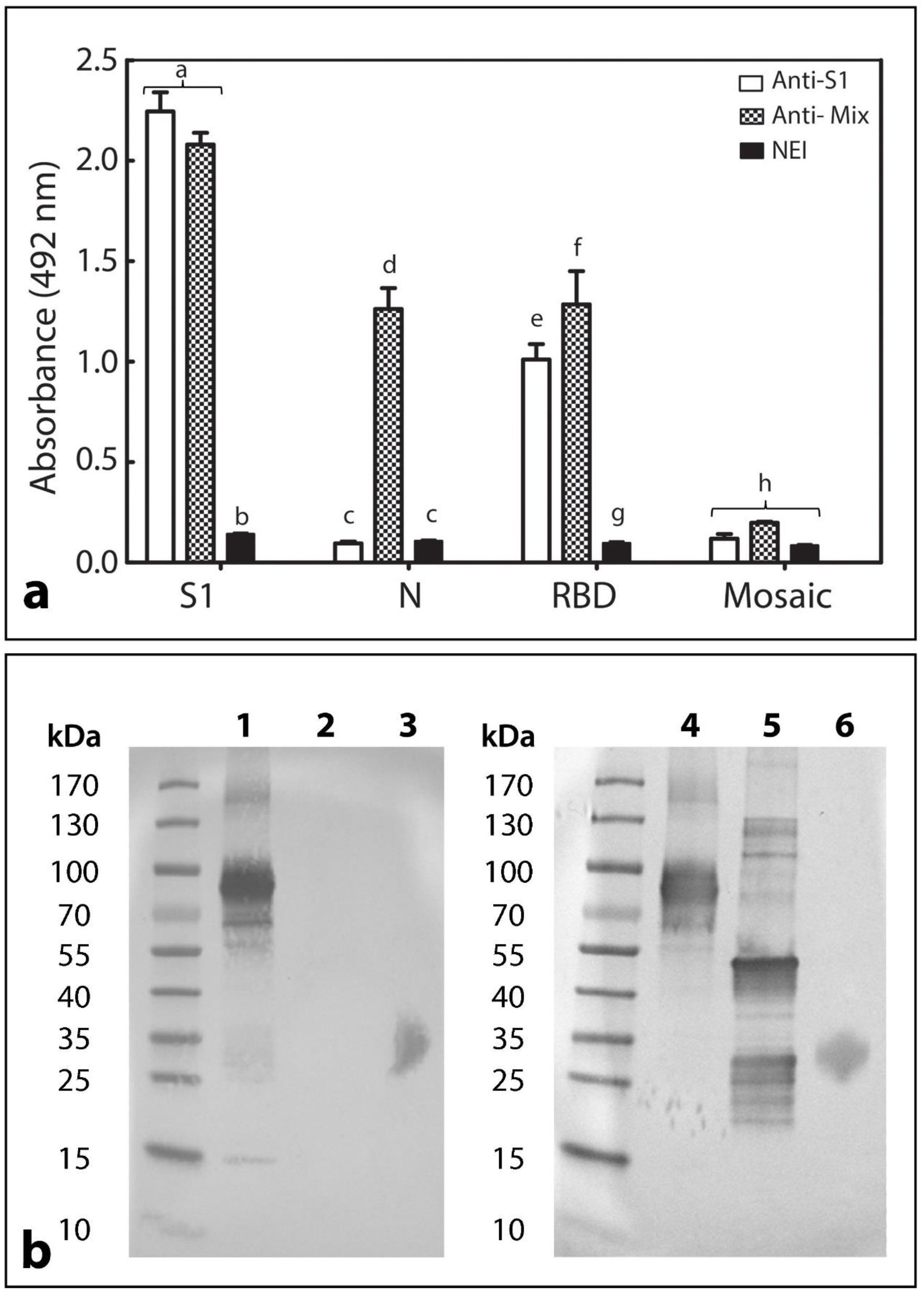
Immunoreactivity profile of anti-S1 and anti-Mix formulations towards SARS- CoV-2 recombinant proteins. a: ELISA antibody response of anti-S1 formulation, anti-Mix formulation, and normal equine immunoglobulin preparation (NEI) towards S1, N, SEM mosaic, and RBD. Samples against S1 and N proteins were assessed at a 1/1000 dilution, and samples against SEM mosaic and RBD were assessed at a 1/500 dilution. Absorbances were recorded at 492 nm. Results are expressed as mean ± SD. A comparison was made between groups by immunogen. Different letters indicate a statistically significant difference (P <0.05). b: Western blot analysis of anti-S1 and anti-Mix formulations towards S1, N, and RBD. Lanes 1-3: immunoreactivity of anti-S1 formulation; Lanes 4-6: immunoreactivity of anti-Mix formulation; Lanes 1 and 4: S1 protein; Lanes 2 and 5: N protein; Lanes 3 and 6: RDB.

The ELISA titers of the final products towards S1 and N was calculated in reference to a normal equine immunoglobulin preparation (NEI) (Table 3). The anti-S1 formulation presented a higher titer against S1 than the anti-Mix formulation, whereas only anti-Mix showed immunoreactivity towards the nucleocapsid viral protein.

The Western blot results (Fig. 4b) agree with the ELISA findings, i.e. anti-S1 formulation immunoreacted with several bands of S1, particularly at 85 kDa band, and with RBD; while anti-Mix formulation immunoreacted with bands of S1, N and RBD proteins. It is worth mentioning that when compared to the electrophoretic profile of viral proteins (Fig. 1), more immunodetected bands appeared in S1 and N, suggesting the presence of other proteins in the recombinant preparations, probably remnants of the upstream process. Because we used proteins that were expressed in non-human cells, there is no risk that immunization could have resulted in the production of equine antibodies towards human proteins that could induce adverse effects.

##### 3.3.3.2 Neutralization profile of final products

We evaluated the ability of the final formulations to neutralize SARS-CoV-2 infection *in vitro* on Vero cells. In general, SARS-CoV-2 inhibition was dose-dependent (Fig. 5). The dilution factor at which the formulations neutralized 50% of the virus (ED50) was 1:29108 (1:26885-1:31643) for anti-S1 and 1:25355 (1:22659-1:58594) for anti-Mix. In previous studies, heterologous formulations towards SARS-CoV-2 RBD were also able to neutralize the infectivity of the virus in cells^25,26^, evidencing that the use of RBD as an immunogen can indeed trigger strong immunoreactivity and neutralization of the virus. Likewise, Cunha and colleagues^27^ showed that immunization of horses with a trimeric spike protein (comprised of both S1 and S2 subunits) generated a formulation with potent *in vitro* neutralizing ability.

**Figure 5.**
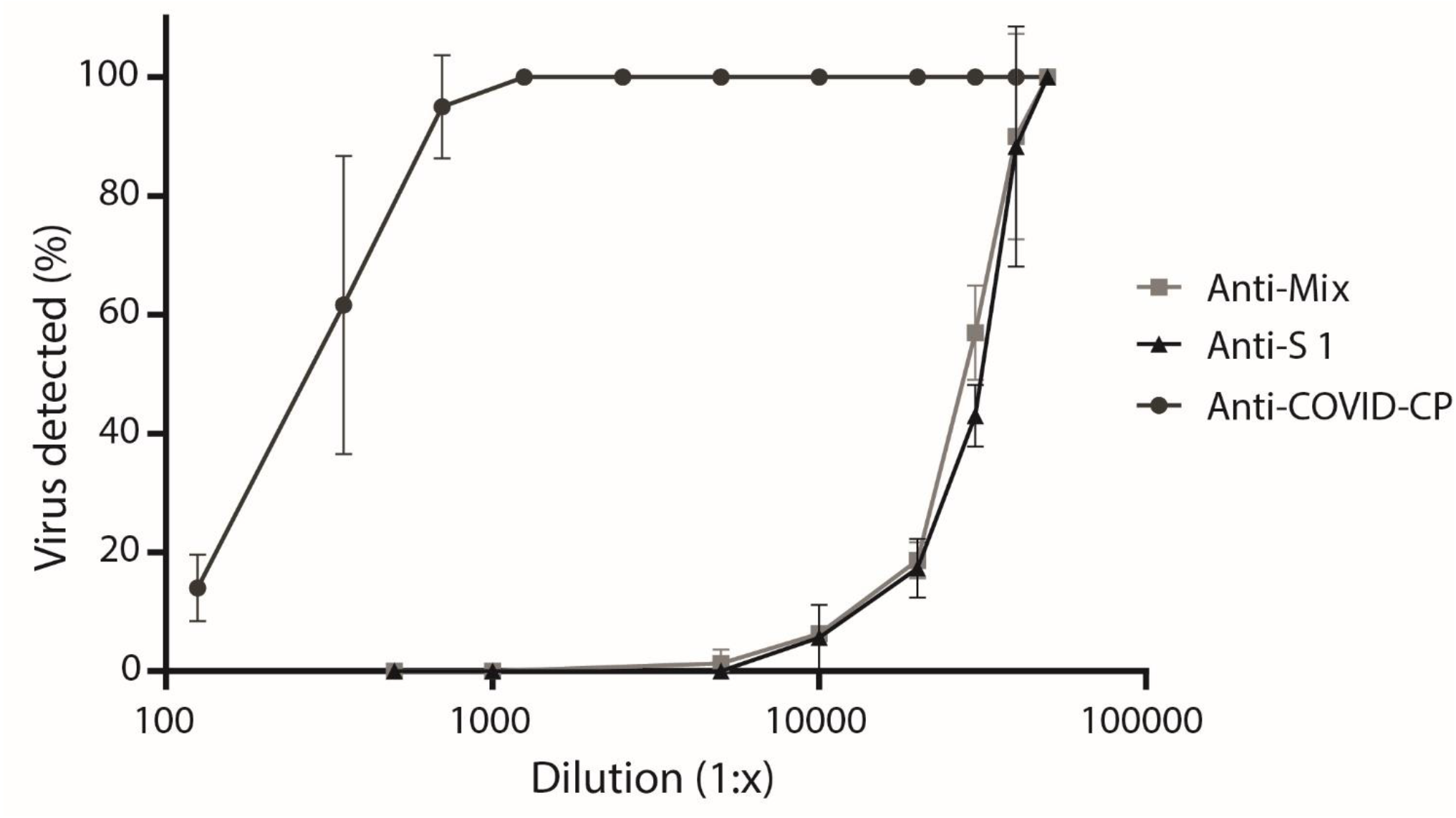
*In vitro* neutralization of SARS-CoV-2 virus by anti-S1 and anti-Mix formulations and human convalescent plasma (Anti-COVID-CP). Samples were serially diluted and their neutralization ability was assessed by a Plaque Reduction Neutralization (PRNT) assay over Vero cells. Results are expressed as mean ± SD of triplicates. ED50 (95 % CI), based on the ability of the preparations to neutralize 50 % of the virus was 1:29108 (1:26885-1:31643), 1:25355 (1:22659-1:58594), and 1:339 (1:295-1:386) for anti-S1 formulation, anti-Mix formulation and human convalescent plasma, respectively.

Here, we demonstrated that immunization of horses with recombinant S1 elicited a strong humoral response. Such a strong neutralizing capacity is most likely due to the presence of anti-RBD antibodies and other antibodies directed towards S protein epitopes such as *the N*-terminal domain (NTD)^44^. During the pre-incubation phase of the PRNT assay with both anti-S1 and anti-Mix formulations, a similar potency of anti-S1 antibodies recognized and bound S1 on the viral surface, neutralizing the infection capacity of the SARS-CoV-2 virus^6^ (Fig. 5).

Given that N protein is an internal antigen, its interaction with specific antibodies in a PRNT assay is unlikely. Thus, we could not assess the contribution of anti-N antibodies to viral neutralization in this study. However, vaccines that have been developed against SARS-CoV and MERS-CoV with targets other than the S protein did not successfully protect animals from infection^10,63^. The contribution of anti-N antibodies to the control of COVID- 19 requires further study.

In terms of the total protein of the formulations required to neutralize 50 % of the viral infection capacity (total protein/ED50), the anti-S1 formulation required 2.3 μg/mL and the anti-Mix formulation required 2.5 μg/mL. Another formulation tested by Pan and colleagues^25^ required 8.8 μg/mL, whereas a formulation reported by Zylberman et al.^26^ reached titer values of 1:10240 with 3 g/dL of total protein. An anti-trimeric S-formulation resulted in a PRNT50 of 1:32000 at 9 g/dL total protein^27^. However, because different methodologies were used to assess the neutralizing ability of the above noted formulations, it would be more appropriate to compare our formulations with convalescent plasma as a control. In this study, ED50 (95% CI) of convalescent plasma was 1:339 (1:295-1:386). In other words, anti-S1 and anti-Mix formulations were 80 times more potent than the convalescent plasma pool (Fig. 5), and a 10-mL vial of equine-derived formulation would be equivalent to 800 mL of human convalescent plasma from donors pre-selected for having a high anti-SARS-CoV-2 titer. These findings are consistent with previous reports^26,27^.

Although the neutralizing potency of the formulations in cell culture is a useful characteristic, it does not necessarily predict clinical efficacy, which also depends on whether antibodies can reach compartments in which the antigen is distributed. Therefore, only properly designed clinical trials can demonstrate the clinical efficacy of heterologous formulations against COVID-19. Once the potency specification of the formulations is established via clinical studies, then cell culture assays can verify consistency and specification compliance within batches in an industrial line.

##### 3.3.3.3 Anti-SARS-CoV-2 equine IgG-mediated activation of human Fcγ receptors (FcγRs)

A surrogate assay was used to evaluate the *in vitro* capacity of equine formulations to trigger the activation of human FcγRIIIA (CD16) (Fig. 6). This assay comprised the cocultivation of anti-viral IgG, previously incubated with S1 and N proteins, with mouse BW:FcγRIIIA-ζ reporter cells expressing the extracellular portion of chimeric human FcγRIIIA. Activation of FcγRIIIA via recognition of the IgG-Fc portion was then determined by measuring IL-2 secretion as a marker^39^. When compared to normal equine IgG (Fig. 6, bar 4), both anti-S1 and anti-Mix formulations induced the secretion of IL-2 (Fig. 6, bars 1 and 2). According to their immunoreactivity, both formulations interacted with S1 protein, and only the anti-Mix formulation reacted to N protein. A pool of anti-SARS-CoV-2 high-titer convalescent human plasma showed significantly higher IL-2 secretion than equine formulations against both S1 and N viral proteins (P < 0.005; Fig. 6, bar 3).

**Figure 6.**
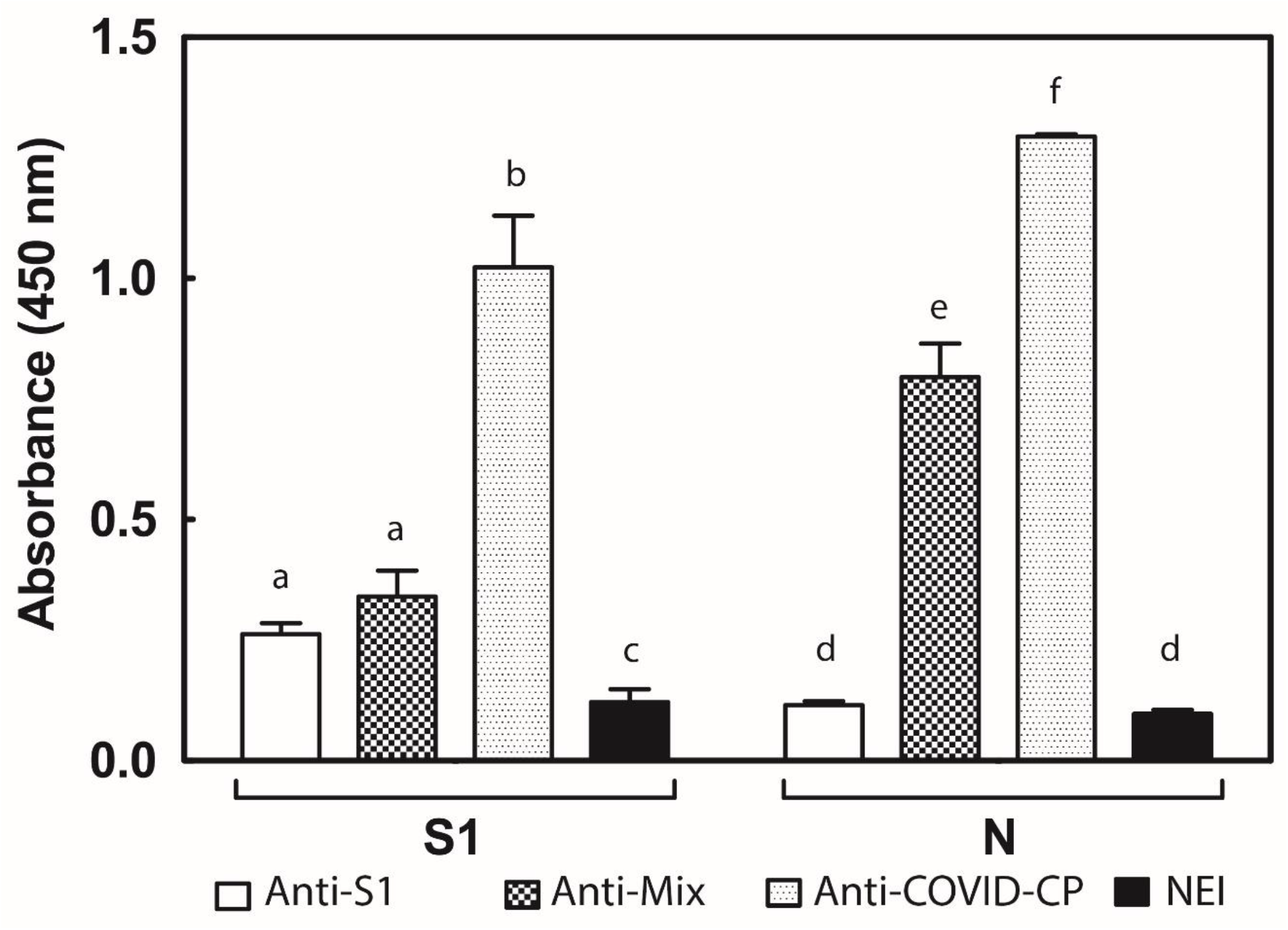
Anti-SARS-CoV-2 equine IgG-mediated activation of human FcγRs. 1: anti-S1 formulation; 2: anti-Mix formulation; 3: anti-SARS-CoV-2 human convalescent plasma pool; 4: Normal equine immunoglobulin preparation (NEI). Plates were coated with 1 μg/well of N and S1 proteins and were incubated with 20 mg/dL of immunoglobulin preparations. BW:FcγRIIIA-ζ transfectants (BW5147 thymoma cells expressing the extracellular portion of human FcγR FcγRIIIA fused to the mouse CD3 ζ-chain) were used. Absorbance was recorded at 450 nm and corresponds to the measurement of mIL-2 by ELISA. Experiments were performed in triplicate and expressed as mean ± SD. A comparison was made between groups by immunogen. Letters indicate statistically significant differences (P <0.05).

Interestingly, our results evidence some capacity of equine IgG to interact and activate human FcγRIIIA *in vitro*, suggesting the possibility of being able also to activate human FcγRs *in vivo.* This finding has several implications in terms of the antiviral mechanism of action and the safety profile of the formulations.

Under this scenario, in addition to neutralizing the viral proteins, anti-viral immunoglobulins may also mediate downstream effector functions via interaction with Fc-receptors in immune cells^10,64^. Such an interaction could lead to the killing of virus-infected cells by effector mechanisms such as antibody-dependent cell-mediated cytotoxicity (ADCC). For example, FcγRIIIA could interact with natural killer cells, or, if equine IgG are able to interact with other human FcγRs, through stimulation of antibody-dependent cellular phagocytosis (ADCP)^65^. Overall, the activation of human FcγRs may represent a clinical advantage of non-digested formulations over F(ab)’2 formulations, although this hypothesis remains to be tested in clinical studies.

At the same time, there are theoretical safety concerns related with the possibility that antiviral hyperimmune immunoglobulin formulations could induce an immune enhancement of the viral disease. Non-neutralizing, low-affinity, or sub-neutralizing concentrations of antibodies that may engage functional responses from human FcγRs, could lead to antibody-dependent enhancement (ADE) or antibody-enhanced immunopathology^8,10,64,65^. Even though ADE and acute lung injury have been reported during SARS-CoV infection in nonhuman models and cell culture studies^66–69^, there is still no compelling clinical, epidemiological or histopathological evidence to support ADE or antibody-enhanced immunopathology of COVID-19 infection or re-infection in humans^10,70^, or during human plasma convalescent therapy^9^. However, both phenomena should be considered as latent risks. Therefore, further studies to elucidate the pathology of SARS-CoV-2 infection as well as strict vigilance during passive immunotherapy with antiviral immunoglobulin formulations are crucial.

In contrast to our protocol, other anti-viral heterologous formulations for the treatment of severe acute respiratory syndromes have included an enzymatic digestion of IgG with pepsin to generate divalent F(ab')2 preparations. Generally, the rationale for Fc removal is to prevent the development of antibody-dependent enhancement (ADE)^20,21,24–27^ and early adverse reactions mediated by the Fc fragment of horse immunoglobulins^20,21^. However, in the particular case of equine antivenoms, similar incidences of early adverse reactions have been reported for F(ab’)2 antivenoms and whole IgG preparations produced by caprylic acid precipitation^59,62^, suggesting that the presence of Fc fragment in the preparation is not the main culprit for these reactions. Additionally, immunoglobulin fragments have shorter half-lives than whole IgG preparations, which confers an advantage to non-digested formulations in terms of active ingredient residence time during administration^42,71^. Clarification of the relative efficacy and safety of F(ab’)2 and IgG anti-SARS-CoV-2 formulations is an important task that must be addressed in the near future.

## Concluding remarks

Taking advantage of our experience with manufacturing snake antivenom, we developed two equine-IgG formulations (anti-S1 and anti-Mix) by immunizing horses with SARS-CoV- 2 recombinant proteins S1, N, and SEM mosaic. Formulations were prepared with a simple, cost-effective, and scalable methodology, and showed high physicochemical and microbiological quality. We demonstrated that horses can produce large quantities of antibodies with high neutralizing potency of the virus *in vitro,* due to the presence of anti-S1 and anti-RBD immunoglobulins in the final products, which demonstrated to be 80 times more potent than a pool of human convalescent plasma. Both formulations have similar pre-clinical quality, safety, and efficacy profiles, but are yet to be validated with proper clinical trials. We suggest that the technological platform presented here could be adapted by other equine immunoglobulin producers worldwide to provide this potential treatment of COVID- 19 in other regions, particularly in low- and middle-income countries.

## Data Availability

All data generated or analyzed during this study are included in this published article.

## Acknowledgements

This work was funded by the Rectoría of the Universidad de Costa Rica, Vicerrectoría de Investigación (Project 741-C0-198) and the Costa Rican National Research Council CONICIT (Project FV-0001-20). The contributions of the Embassy of the People’s Republic of China in Costa Rica and Roche Servicios S.A. (Costa Rica), as well as the strong support of the Rector of UCR, Dr. Carlos Araya, and the Costa Rican President, Carlos Alvarado, are greatly appreciated. Support by way of laboratory reagents and analysis by Diagnóstico Albéitar, Tecnodiagnóstica, and Biocientífica International are also acknowledged. The group thanks Melania Alfaro, Sergio Rojas, Rodolfo Cruz, Flory Cruz, Omar González, Julio Lara, Roberto Federspiel, and Gabriela Quirce for their generous donation of horses, and Jenny Stynoski for the detailed review of English and writing.

## Author contributions statement

G.L., M.H, M.Va., C.D. and A.A conceived the work and designed the experiments.

M.H., M.Va., M.A., A. Sa., A.Se., G.S., E.C.-A., K.R., A.N., C.B., M.Vi., Ad.Sa., D.C., D.S., G.D., E.S., M.C., D.U., E.M., R.E., J.G., M.M. and A.C.C performed the experiments.

G.L., M.H., M.Va., M.A., A. Sa., A.Se., A.G., E.C.-A., K.R., A.N., C.B., M.Vi., A.H, C.D. and A.A. analyzed and interpreted the data.

G.L., M.H, M.Va, M.A. and A.G. wrote the manuscript.

A.Se., A.G., G.S., E.C.-A, M.Vi., L.S., J.M.G and C.D. substantively reviewed the manuscript.

