## Supplementary material for "Development and pre-clinical characterization of two therapeutic equine formulations towards SARS-CoV-2 proteins for the potential treatment of COVID-19": Figure S1

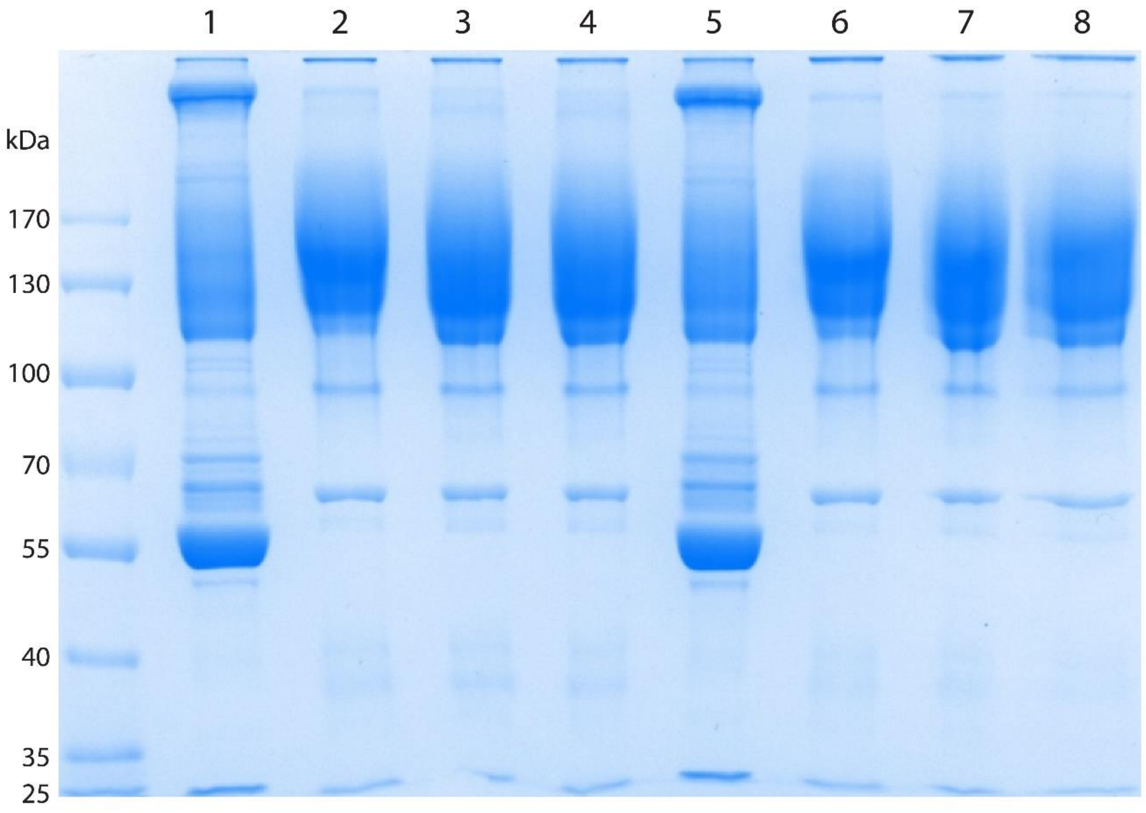


**Figure S1**. SDS-PAGE analysis of pools of plasma and final anti-S1 and anti-Mix formulations. Lane 1: Hyperimmune pool of plasma anti-S1; Lanes 2-4: Anti-S1 formulation; Lane53: Hyperimmune pool of plasma anti-Mix; Lanes 6-8: Anti-Mix formulation. Samples (20 µg) were loaded in a 7.5% polyacrylamide gel in the presence of SDS and run under non-reducing conditions. The gel was stained with Coomassie Blue.
